# GMQN: A reference-based method for correcting batch effects as well as probes bias in HumanMethylation BeadChip

**DOI:** 10.1101/2021.09.06.459116

**Authors:** Zhuang Xiong, Mengwei Li, Yingke Ma, Rujiao Li, Yiming Bao

## Abstract

Illumina HumanMethylation BeadChip is one of the most cost-effective ways to quantify DNA methylation levels at the single-base level across the human genome, which makes it a routine platform for epigenome-wide association studies. It has accumulated tens of thousands of DNA methylation array samples in public databases, thus provide great support for data integration and further analysis. However, majority of public DNA methylation data are deposited as processed data without background probes which are widely used in data normalization. Here we present Gaussian mixture quantile normalization (GMQN), a reference based method for correcting batch effects as well as probes bias in HumanMethylation BeadChip. Availability and implementation: https://github.com/MengweiLi-project/gmqn.

## 1 Introduction

As a well-known epigenetic marker, DNA methylation plays a crucial role in numerous physiological processes and also complex traits, such as development, phenotype and cancer[1-3]. With the advancement of epigenetic sequencing technologies and the reduction of sequencing costs, especially the DNA methylation array, massive samples can be used to explore epigenetic basis of complex traits, which has also resulted in the accumulation of a large number of DNA methylation array data in public databases[4-11]. According to the statistics of DNA methylation array data in the GEO database, among them, Illumina HumanMethylation450 BeadChip (450k) has become the most widely used means of large-scale methylation profiling of human samples in recent years. The newly emerging Illumina HumanMethylationEPIC BeadChip (EPIC/850k) uses the same technology as 450k, but covers at nearly double the number of CpG sites, and will become the main effective strategy of epigenome-wide association studies (EWAS) in the future (figure 1A). Integrating both large samples from public resources and private data will become a common and main research strategy for future research on potential regulatory mechanisms of complex traits, particularly for EWAS[12]. Different laboratories have differences in sample processing and sequencing processes. These differences have nothing to do with biological factors and are defined as between-array bias (batch effects) [13, 14], which will reduce the signal-to-noise ratio and adversely affect downstream analysis in the integration of public data.

**Figure 1a:**
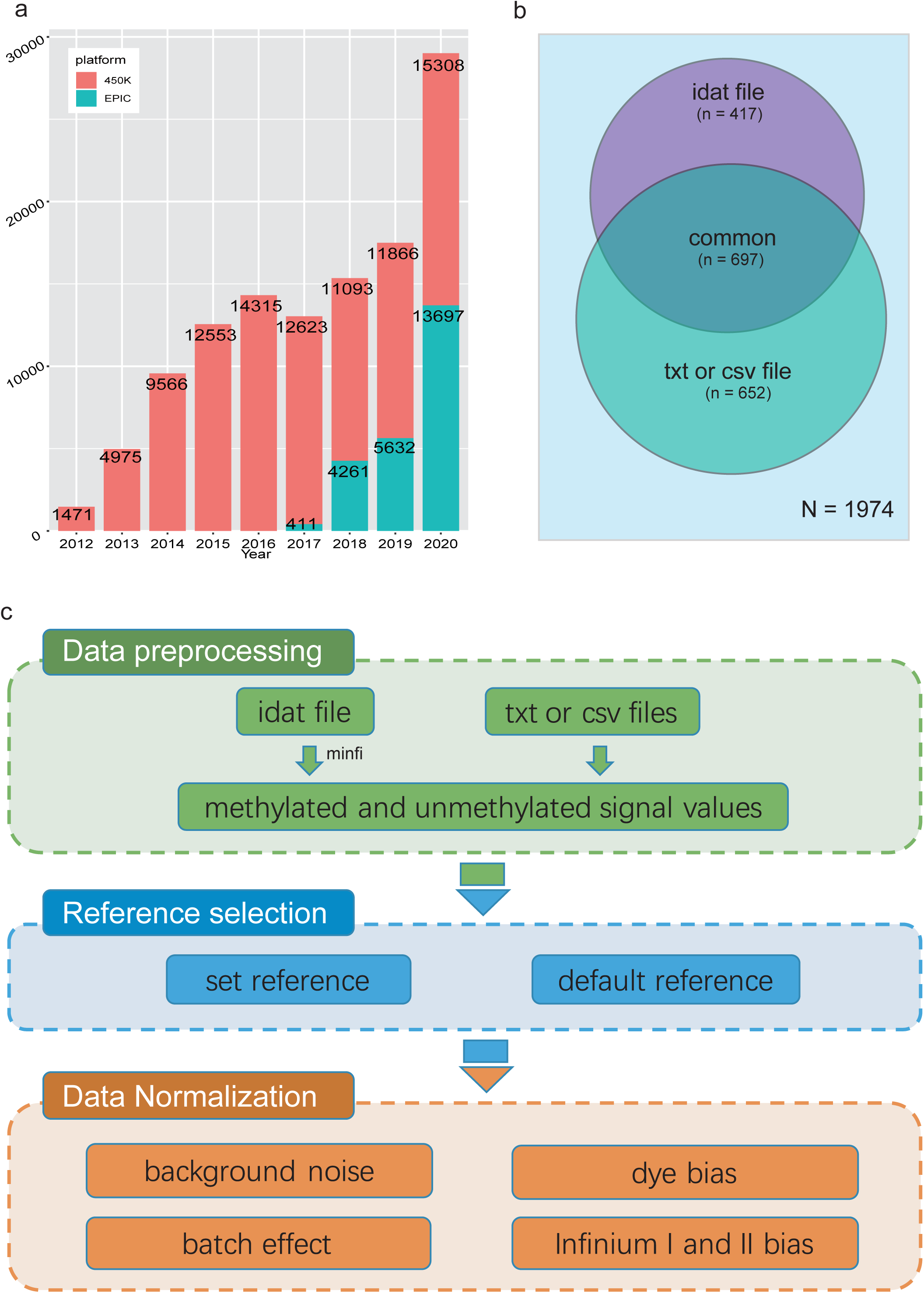
Statistics of 450k and EPIC data by year and project number in NCBI GEO database. b: Distribution of data types of DNA methylation chip projects submitted in the GEO database as of December 2020. There are a total of 1,114 items containing the original ‘idat’ files, and 1,349 items containing TXT or CSV files, indicating that most of the items miss original file. c: The workflow of GMQN.

Currently, a number of DNA methylation array normalization methods have been developed, each with its own set of advantages and disadvantages in different study scenarios. Many methods, on the other hand, are not well suited to the analysis of large amount of public data. The majority of methods depend on data from control probes or OOB (Out of Band) probes, and they can’t be used for public data unless the original data is available. However, only about half of the 450k and EPIC projects in GEO, the largest publicly accessible DNA methylation array database, provide original data (figure 1B). As the well-known normalization method on *β*-values of DNA methylation, SWAN and BMIQ do not use the information from these two types of probes, instead, they only deal with within-array bias. (Infinium I and II bias) [15, 16].

Without violating these constraints, we must still deal with four types of deviations: Infinium I and II bias, red and green signal deviations, background noise, and batch effects. Therefore, we propose a reference-based method for correcting batch effects as well as probes bias in HumanMethylation BeadChip, which is called Gaussian Mixture Quantile Normalization (GMQN). The method includes four steps: (I) Fitting of a two-state Gaussian mixture model to the median values of each Infinium I probe signal intensity from a large single study (GSE105018). The mean and variance of two components are used as reference for rescaling Infinium I probes. (II) Fitting of a two-state Gaussian mixture model to the input Infinium I probe signal intensity. (III) For Infinium I probes from each component of input data, transform their probabilities to quantiles using the inverse of the cumulative Gaussian distribution with mean and variance estimated from the corresponding reference component. (IV) After the batch effect is reversed, GMQN can also normalize Infinium II probes on the basis of Infinium I probes in combination with BMIQ and SWAN, the two well-known normalization method on β-values of DNA methylation[15, 16](figure 1C).

## 2 Materials and Methods

### 2.1 DNA methylation data

The data for method development and testing are taken from the GEO and TCGA databases, which contain 450k and EPIC records (Table 1). Respectively, the sample information is annotated using a combination of automated grabbing and manual analysis. The R package “minfi” (http://www.bioconductor.org/packages/release/bioc/html/minfi.html) is primarily used to interpret and preprocess the original signal[17]. Considering that some public data only have original methylated and unmethylated signal value files, we use the “preprocessRaw” method to extract the original signal values without any processing. To ensure fairness, the methylated and unmethylated signal values of all probes except the control probe are collected and used as the input value in all subsequent tests and comparisons. The methylation level is represented by *β, β* = *M*/(*M* + *U*), where *M* and *U* represent the intensity of methylation and non-methylation signal values, respectively.

**Table1.** Overview of benchmark test dataset

| Project id | Number of samples | Benchmark Test | Annotation | platform |
| --- | --- | --- | --- | --- |
| GSE52731 | 56 | batch effects detection | - | 450k |
| GSE139687 | 27 | batch effects detection | - | EPIC |
| GSE42861 | 689 | case-control study | Rheumatoid Arthritis | 450k |
| GSE128235 | 537 | case-control study | Depression | 450k |
| GSE125105 | 210 | regression analysis | Age | 450k |
| GSE42861 | 335 | regression analysis | Age | 450k |
| GSE87571 | 732 | regression analysis | Age | 450k |
| GSE87571 | 732 | comparison of the methylation levels of adjacent sites | - | 450k |

### 2.2 Reference data

In GMQN, there are two ways to set the reference signal value distribution. To begin, users can use the function ‘set reference’ in the ‘GMQN’ package to match their own data to fit their own reference distribution. The second option is to use the default reference, which is a two-state Gaussian mixture model fitted to the median values of each Infinium I probe signal intensity from a large single study (GSE105018). The mean and variance of two components are used as reference for rescaling Infinium I probes. This project includes 1658 whole blood samples obtained from E-Risk cohort participants when they were 18 years old[18].

### 2.3 GMQN

In order to remove any source of variation that is not related to biology but rather to technical limitations, such as dye bias or batch effect, we first need to find the manifestations of these variations in the data[19]. To that end, we investigated the signal value distribution characteristics of two types of probes. We found that the signal values of the red and green channels of Infinium I probes can be decomposed into the superposition of two Gaussian distributions, and that the fitting parameters of these Gaussian distributions can effectively discern batches (details in result). Using this feature, we draw on the idea of BMIQ, respectively fit the Gaussian mixture distribution to the signal values of the red and green channels of Infinium I probes, and then adjust the shape of the Gaussian distribution corresponding to different samples to the same shape to the reference to minimize batch effects and other deviations. To achieve this process, GMQN standardizes the data in three steps.

The first step is the establishment of the reference distribution. In order to address the issue in rapid growth of public data, GMQN adopts a data normalization method based on reference distribution, which is also widely used in the normalization of data in the EWAS Data Hub database[4]. Usually, we need to average the signal intensity of each probe on the reference data set between samples, and fit the Gaussian mixture distribution to the probe signal intensity on the red and green channels of Infinium I probes respectively. The Expectation-Maximization algorithm is used to estimate the parameters, and the red channel fitting result is expressed as: 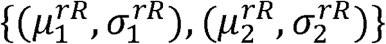, the green channel fitting result is expressed as: 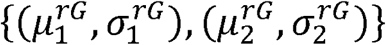, where *r* is the reference, 1 and 2 respectively represent the two states of the mixed model with a smaller and larger mean, and *R* and *G* represent the red and green channels, respectively.

The second step is the normalization between arrays. The between-array normalization is carried out separately for the red and green channels of Infinium I probes. Taking the green channel as an example, we first fit the Gaussian mixture distribution to the signal intensity of the green channel of the input Infinium I probe to obtain the fitting parameters 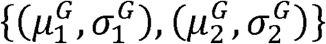. For the state with the smaller mean value, state 1, we perform the following conversion:

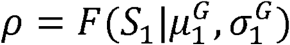

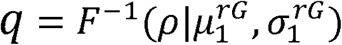

Where *S*_1_ is the signal belonging to state 1 in the green signal, *ρ* is the cumulative distribution probability of the signal value in the Gaussian distribution, and *q* is the signal value corresponding to the cumulative probability in the reference distribution. Through this step, we map the input signal to the reference signal, eliminate biases such as dye bias and batch effect. The signal in state 2 and the signal in the red channel are processed using similar steps.

The third step is within-array normalization, which mainly including Infinium I/II-type bias correction. In the second step, we obtained the normalized Infinium I probes signal. Based on Infinium I probes signal, we use BMIQ or SWAN to standardize the Infinium II probes signal. We respectively made some fine-tunings to BMIQ and SWAN to improve speed and effectiveness.

### 2.4 Benchmark Test

Since other methods cannot be used in the absence of original data, in the benchmark test, we compare GMQN, SWAN, BMIQ, and GMQN combined with SWAN and BMIQ. In order to test whether GMQN can improve the effect of SWAN and BMIQ, we designed the following four benchmark tests.

#### 2.4.1 Batch effects detection

In order to make the method more universal, we searched the GEO database for two sets of technical replicates, including 450k and 850k chips. The first group (EPIC, GSE139687) has 9 samples that are replicated three times each, while the second group (450k, GSE52731) has 56 repetitions of one sample. For the first set of samples, we measured the variance at the probe level between every three technical replicates and then averaged the variance among the nine samples. For the second set of samples, we directly calculate the variance of the sample at the probe level.

#### 2.4.2 Case-control study

Case-control study is the most common form of research in EWAS. Researchers classify samples into case and control groups and look for differences in methylation sites between the two groups in this form of study. We used the data of two diseases in public databases in order to assess the performance of GMQN in the case-control studies. To simulate two separate batches, we divide the samples in the data set 2:1 into training and test sets for each disease. In the training set, we aim to keep the samples in the same batch of chips, and the batch effect and other errors are kept to a minimum. Differential methylation analysis is carried out in both the training and test sets, with the training set’s findings serving as the gold standard for detecting consistency between the training and test sets and drawing the receiver operating characteristic (ROC) curve.

#### 2.4.3 Regression analysis

The term “regression analysis” refers to the process of associating DNA methylation levels with continuous variables such as age, BMI, and so on in order to identify DNA methylation sites that are linked to these variables. Age is a trait that has been reported more frequently in EWAS, and there is a lot of data on it. As a result, we used age as the research object in this study and obtained 1277 sample data sets containing age information from three separate projects. The data from multiple items ensures that the sample’s batch effect is high, allowing each standardized method’s effect to be better measured. A large number of studies have reported that there is a linear relationship between DNA methylation and age[20, 21], and the Pearson Correlation Coefficient is particularly suitable for the quantification of the linear relationship. Therefore, we calculated the Pearson correlation coefficient between DNA methylation and age as quantitative indicators.

#### 2.4.4 Comparison of the methylation levels of adjacent CpG sites

Studies have reported that DNA methyltransferase has a limited range of action, resulting in nearly identical methylation levels at adjacent CpG sites on the genome[22, 23]. In this part, we selected 141,653 pairs of probes with a genome distance of less than 10 bps on the chip. For each sample, we calculated the average of the difference in DNA methylation levels of these probes, and chose 141,653 pairs of probes at random as controls.

## 3 Results

### 3.1 The signal intensity distribution characteristics of Infinium I probes and the principle of GMQN

The signal from the control probe can, ideally, be used to quantify the batch effect between samples. However, most public data lack original data, so we tried to find other manifestations of batch effects. We found that the signal intensity of the red and green channels of Infinium I probes can be approximately decomposed into the superposition of two Gaussian distributions, both in 450k and EPIC array (Figure 2). We speculate that this may be related to the bimodal distribution of human DNA methylation level. When the methylation value is extremely high (>0.8) or extremely low (<0.2), one of the two Infinium I probes that detects the site’s methylation level emits almost no light, and the fluorescence signal intensity of these probes constitutes the first peak of the Gaussian distribution, that is, the peak with the smaller mean. The fluorescence signal intensity of other probes constitutes the second Gaussian distribution. Since the methylation levels of the sites corresponding to these probes are dispersed, the Gaussian distribution’s variance is larger. We cluster the Gaussian distribution parameters fitted by different samples to see if these Gaussian peaks are related to batches. The results show that the fitting parameters of the four Gaussian distributions (two for each of the red and green channels) can be used to differentiate the batches, and that even if the sample difference is large, the parameter difference will be small within the batches (Figure 2).

**Figure 2:**
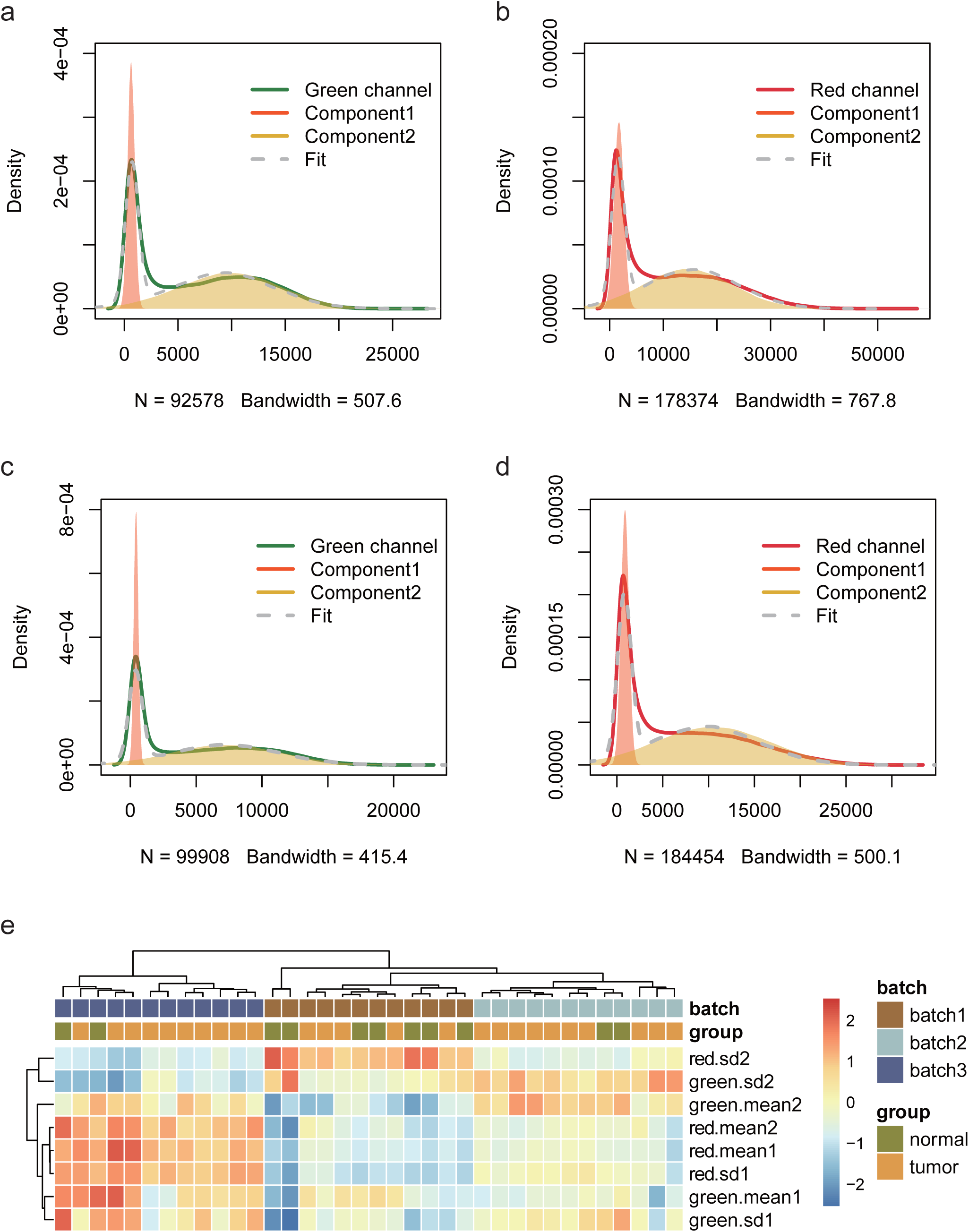
The signal intensity distribution characteristics of Infinium I probes (450k data (a, b), EPIC data (c, d)) and clustering results of different batches of samples based on fitting parameters of the Gaussian distributions (e).

Using this feature, we propose a GMQN standardization method. The basic principle of this method is to fit a Gaussian mixture model for Infinium I probes of different batches, and then adjust the Gaussian distribution shapes fitted by different batches to the same to eliminate the batch effect on Infinium I probes. Finally, take the Infinium I probes as the standard, and use BMIQ or SWAN to standardize the Infinium II probes. The signal strength distribution of the red and green channels of Infinium I probes was then measured in two batches of samples in a TCGA tumor project before and after GMQN normalization. We found that the distribution of the two batches differs greatly both in 450k and EPIC data, and the differences are not due to biological differences (tumor and normal). The distributions of the two batches tend to be consistent after GMQN standardization (Figure 3, Figure 3S).

**Figure 3:**
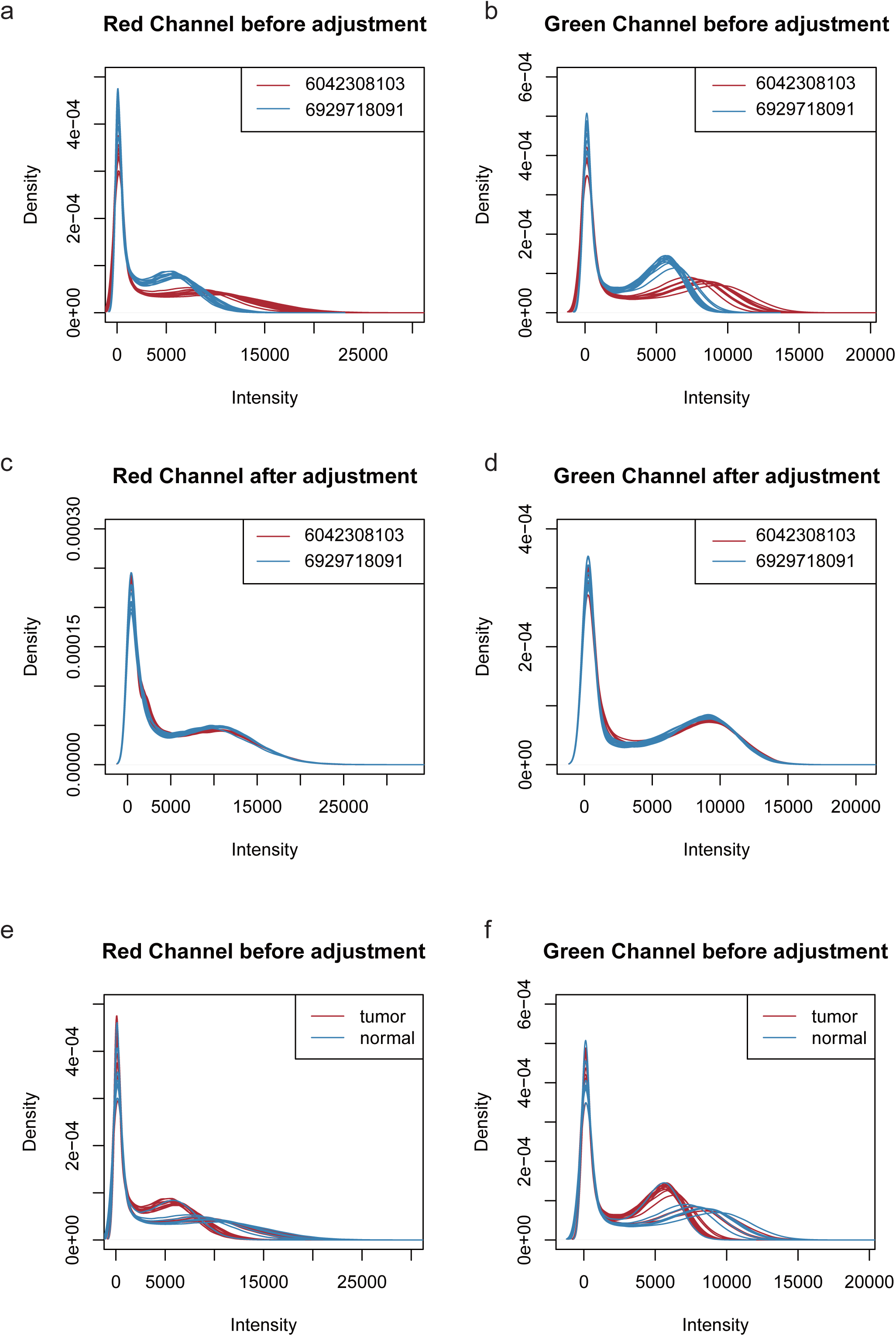
The 450k data signal strength distribution of the red and green channels of Infinium I probes before and after GMQN normalization. The signal intensities of the red (a) and green (b) channels of the two batches were clearly divided into two batches before being corrected. And the differences are not due to biological differences (tumor and normal) (e, f). After the GMQN correction, the batch effect problem is significantly reduced (c, d).

### 3.2 GMQN reduces technical variability

Technical repetition is the most direct way to measure the batch effect. As a result, we chose two different sets of technical replicates. The first set (EPIC, GSE139687) has 9 samples that are repeated three times each, while the second set (450k, GSE52731) has 56 repetitions of one sample[24, 25]. The variances of the probes of the two sets of samples were determined separately. While each method decreases the variance of the probe methylation level relative to the original data in the two sets of technical replicates, the variance of the probe methylation level after GMQN+BMIQ and GMQN+SWAN treatment is the lowest and second lowest, respectively (Figure 4a, 4b). In particular, without combining SWAN and BMIQ, GMQN performed best in the first data set (Figure 4a). This demonstrates that GMQN, especially used in combination with BMIQ and SWAN, is capable of effectively reducing batch effects.

**Figure 4:**
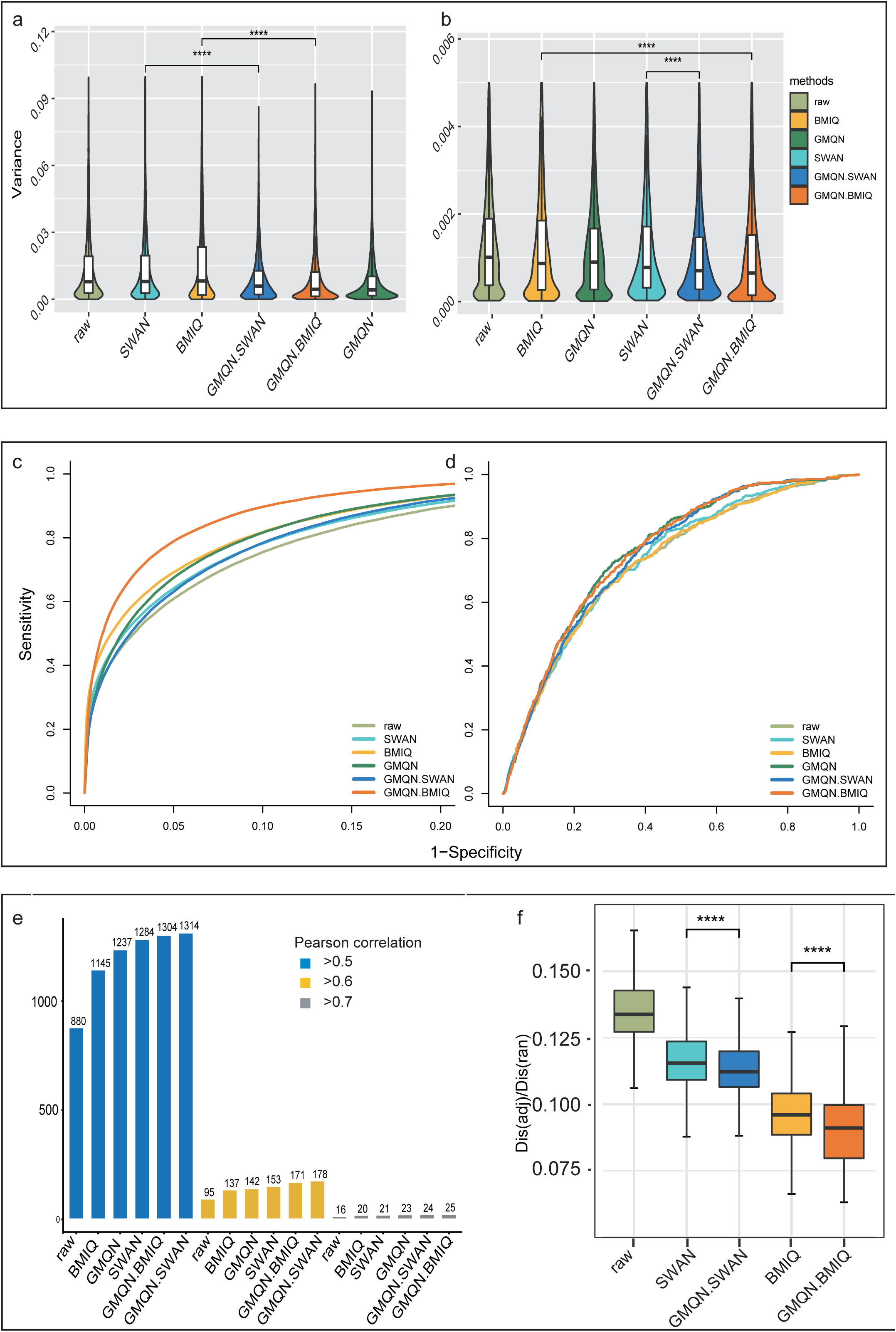
The result of Benchmark Test. a and b: batch effects detection c and d: case-control study e: regression analysis f: comparison of the methylation levels of adjacent CpG sites (**** p<10-4, **** p<10-4)

### 3.3 GMQN leads to better detection of differential methylation

In order to test the effects of GMQN in the case-control studies, we selected normal and disease samples for rheumatoid arthritis and depression[26, 27]. The differential methylation estimation results indicate that there are approximately 50,000 and 1,000 differential methylation positions in the normal and disease samples of these two diseases, respectively. The ROC curve shows that compared with the original data, the consistency of the training set and the test set results is greatly improved in rheumatoid arthritis, GMQN + BMIQ has the best effect, while SWAN and the original data have poor results, but whether it is BMIQ or SWAN, the effect can be achieved after combination with GMQN, GMQN, GMQN+BMIQ, and GMQN+SWAN all outperform other methods in the depression group (Figure 4c, 4d). In case-control studies, these results suggest that GMQN can enhance SWAN and BMIQ effects.

### 3.4 GMQN improves the effectiveness of regression analysis

Regression analysis is a crucial form of analysis in EWAS. For continuous traits such as age and BMI, the relevant DNA methylation sites can be found through regression analysis. Compared with case-control study, the results of regression analysis are often more influenced by data processing methods.

We used data processed by different methods to identify age-related DNA methylation sites to examine the effect of GMQN in regression analysis. Our data in this analysis comes from three separate projects, the batch effect is high, and the sample age period is also large (from 14 years old to 94 years old)[25, 26, 28]. Using the Pearson correlation coefficients of 0.5, 0.6, and 0.7 as thresholds, we measured the number of age-related DNA methylation sites identified by each method. The findings show that the GMQN+SWAN treatment group can find more age-related methylation sites than other methods under various thresholds, and GMQN can boost the effects of BMIQ and SWAN under a strict threshold, and improve the effect of regression analysis (Figure4e). In order to ensure that the sites found by GMQN are true positive sites, we further analyze these sites. Surprisingly, we examined the 5 sites (cg15448975, cg16419235, cg07416237, cg04875128, cg14692377) with Pearson correlation coefficient less than 0.7 after BMIQ analysis and greater than 0.7 after GMQN+BMIQ analysis in the EWAS Atlas, a curated knowledgebase of epigenome-wide association studies[29], and discovered all of them were age-related, indicating that the majority of the newly discovered age-related sites in GMQN are true positives.

### 3.5 GMQN reduces differences in methylation levels between adjacent CpG sites

The difference in methylation levels between adjacent CpG sites is approximately 12% of the difference between random sites. Simultaneously, the difference in methylation levels between adjacent CpG sites in the original data group is greater than in other groups, confirming that this benchmark test is reasonable. The GMQN+BMIQ processed group had the smallest difference in methylation levels between adjacent CpG sites, while the GMQN+SWAN treatment was not as efficient as BMIQ but still better than SWAN (Figure 4f).

## 4 Discussion

The accumulation of public DNA methylation array data has provided favorable conditions for the advancement of EWAS, allowing data analysts to investigate the relationship between various traits by massive public data mining without relying on experiments. As a result, we developed GMQN, a standardized method suitable for massive public DNA methylation array data. GMQN has the following benefits over other methylation chip data standardization methods: First and foremost, GMQN is a reference-based Gaussian mixture quantile normalization method. It can be used to calibrate a newly added sample to the same level as the previous batch of samples without wasting a lot of computational resources, which will solve the N+1 issue in big data integration. The EWAS Data Hub database currently integrates and stores 95,738 methylation chip data using the GMQN[4]. Secondly, GMQN will address the issue of batch effect processing and standardization in public data due to missing original data, making it easier for researchers to combine self-produced and public data to investigate epigenetic mechanisms of various phenotypes. Finally, since most DNA methylation chip processing software packages are written in R, GMQN is written in R as well to increase compatibility with other software. Users can easily achieve GMQN standardization using the R package “GMQN.” Users can combine SWAN and BMIQ to perform parallel analysis on multiple CPUs using the two functions “gmqn_swan_parallel” and “gmqn_bmiq_parallel”.

By evaluating 450k and EPIC array data in four separate application scenarios above, we found that GMQN can effectively minimize noise in public data and increase the accuracy of downstream analysis. GMQN will boost the two well-known methylation chip standardization methods, BMIQ and SWAN, even if it does not perform well in some scenarios, especially when the reference methylation distribution and the methylation data distribution to be standardized are vastly different, as in DNA methyltransferase gene knockout samples versus normal samples. Many DNA methylation array data standardization methods have been developed in recent years[30-36], and they have proven to be invaluable in epigenetics research, especially for EWAS[37, 38]. However, we believe that GMQN can improve the normalization effect to some degree, especially when there is no original data.

## Acknowledgement

Strategic Priority Research Program of the Chinese Academy of Sciences [XDB38030200, XDB38030400]; Key Technology Talent Program of the Chinese Academy of Sciences

---

Figure 3S: The EPIC data signal strength distribution of the red and green channels of Infinium I probes before and after GMQN normalization.

