## Supplementary figures and images for "GMQN: A reference-based method for correcting batch effects as well as probes bias in HumanMethylation BeadChip"

### Supplemental Figure3S

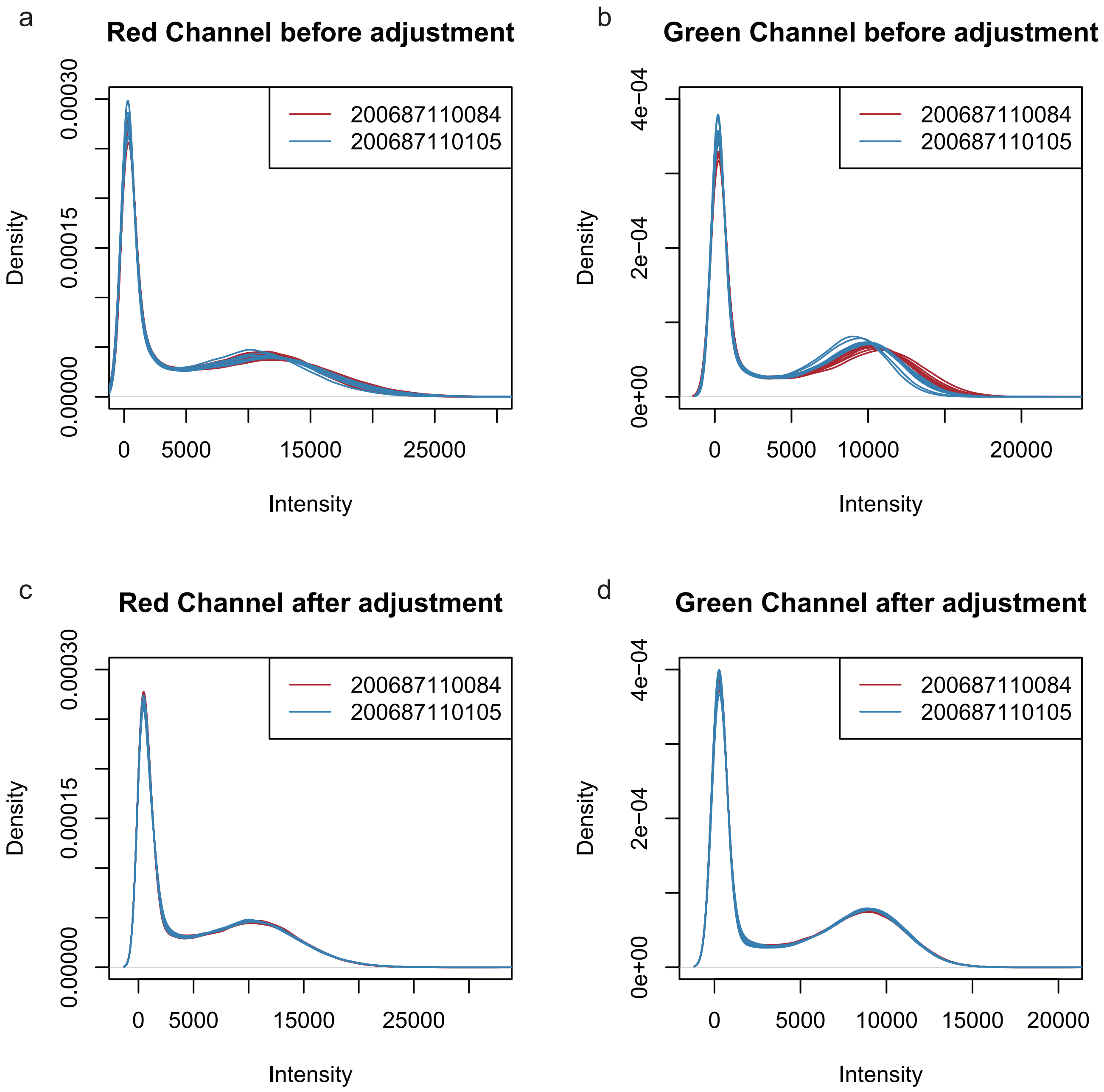
